## Supplementary material for "Optogenetic Manipulation of Cell Migration with High Spatiotemporal Resolution Using Lattice Lightsheet Microscopy": https://drive.google.com/file/d/1GUEtgMkFocTPm6ubJPR9R01jglei6UK5/view?usp=sharing: MS_optoLLSM_Supp.pdf

### Comparison of Bessel lightsheet and Gaussian lightsheet created by objectives with various NAs

For the optogenetic experiment, the stimulation beam is used to activate the optogenetic molecules in cells. In principle, the stimulation beam can activate molecules in the beam path (except for two-photon stimulation). However, the detection schemes used in the experiment may create misleading images where all fluorescence signals along the propagation direction are projected onto a 2D image. To compare different excitation and detection schemes, the beam paths in the samples with different excitation schemes (wide-field, confocal, Gaussian lightsheet, and Bessel lightsheet) are depicted in Fig. S1. In figure S1a, a wide field excitation with a low NA objective is used where the fluorescence signals are detected by the same objective. With a high NA objective, a tightly focused beam is formed, as shown in figure S1b, where a pinhole in front of the detector can reject the out-of-focus signals forming confocal images. In both cases, the excitation beams activate molecules along the beam path, and the signals are projected onto a 2D image. For lightsheet microscopy, different objectives are used for excitation and detection. Shown in figure S1c and S1d are Gaussian and Bessel lightsheet illumination, respectively, where a separate detection objective at the right angle can image all the activated molecules along the beam propagation direction. To characterize the Bessel beam used in this experiment, we simulated and measured Bessel beam profiles in XY and XZ directions, as shown in figure S2. The point-spread function (PSF) along the beam propagation direction is calculated and measured at three spots marked in a white circle with a cross. In this experiment, maximum/outer and minimum/inner numerical apertures (NA) were 0.64 and 0.56, respectively. From the simulated and measured beam profiles, we can find that there are some intensity contributions from the side lobes. To compare with the light sheet created by the Gaussian beam, we need to quantify the intensity contribution from the main lobe and side lobes. We calculated the beam length, thickness, and effective intensity of the Bessel beam with various inner NAs at a fixed outer NA (=0.64). Shown in figure S9 is the beam profile of a Bessel beam near its focal point. The beam propagates along the $y$ axis. We assume that the detection lens is located at the $+z$ side. Therefore, the field distribution on the $xy$ plane is imaged. As indicated in Fig. S2, a Bessel beam is composed of the main lobe and many side lobes. The side of each lobe can be characterized by the full width at half maxima (FWHMs) along with the lateral and propagation directions, which are denoted as $\mathrm{FWHM}_{x,n}$ (thickness) and $\mathrm{FWHM}_{y,n}$ (length), respectively, where $n$ represents $\mathrm{n^{th}}$ side lobe in the Bessel beam. Fig. S9b and S9c are the calculated $\mathrm{FWHM}_{x,n}$ and $\mathrm{FWHM}_{y,n}$ using the fast Fourier transformation under different sizes of ring apertures. The wavelength is 0.488 μm. The outer numerical aperture $\mathrm{NA_{out}}$ defined as $\arctan(R_{\mathrm{out}}/f)$ is set to 0.64, where $R_{\mathrm{out}}$ is the outer radius of the aperture, and $f$ is the focal length. We calculate the thickness, length, and effective intensity of the main lobe and each side lobe at a different inner numerical aperture $\mathrm{NA_{in}} \equiv \arctan(R_{\mathrm{in}}/f)$, where $R_{\mathrm{in}}$ is the inner radius of the aperture. For both the main and side lobes, the thicknesses ($\mathrm{FWHM}_{x,n}$ ) are only reduced slightly as $\mathrm{NA_{in}}$ increases (thinner ring aperture) as shown in figure S9b. This is because $\mathrm{FWHM}_{x,n}$ of the main and side lobes decreases with the average radius $(R_{\mathrm{out}} + R_{\mathrm{in}})/2$ of aperture, which is only varied slightly by $R_{\mathrm{in}}$ in our cases. The thickness of the main lobe is also larger than those of the side lobes, which is a feature of the zero$^{\mathrm{th}}$-order Bessel function. On the other hand, the lengths of all the other side lobes ($\mathrm{FWHM}_{y,n}$) increase significantly with $\mathrm{NA_{in}}$ (reduced aperture opening) due to the uncertainty principle as shown in Fig. S9c. Typically, the length of side lobes is prolonged more than the main lobe as $\mathrm{NA_{in}}$ increases, but their increment ratios do not differ much. To calculate the effective intensity $I_n$ inside a rectangular region $\Omega_n$ defined by $\mathrm{FWHM}_{x,n}$ and $\mathrm{FWHM}_{y,n}$, we set the input power $at$ $P_{\mathrm{in}} = 1$ μW at the ring aperture and calculate the effective intensity using the following equation:

$$I_n \equiv \frac{\frac{n_{\mathrm{a}}}{2\eta_0}\int_{\Omega_n} \mathrm{d}\boldsymbol{\rho}|\mathbf{E}(\boldsymbol{\rho})|^2}{\mathrm{FWHM}_{x,n}\times\mathrm{FWHM}_{y,n}}$$

where $n_{\mathrm{a}} = 1.33$ is the refractive index of water (ambiance); $\eta_0 = 377\ \Omega$ is the intrinsic impedance. As shown in Fig. S9d, the effective intensity $I_n$ drops $as$ $\mathrm{NA_{in}}$ increases as a result of the prolonged beam. In addition, the effective intensity of the main lobe is much higher than all other side lobes indicating that intensity contribution from side lobes of Bessel beams can be neglected in the Bessel lightsheet microscopy,

For comparison, we calculate the same beam properties for the Gaussian beam. As illustrated in Fig. S10a, the thickness, length, and effective energy of the Gaussian beam are calculated as a function of the numerical aperture $\mathrm{NA} \equiv \arctan(W/f)$ of Gaussian beam, where $W$ is the beam waist behind the excitation lens. As shown in Fig. S10b, unlike the thickness ( $\mathrm{FWHM}_x$ ) of the Bessel beam, the thickness of the Gaussian beam significantly decreases toward the diffraction limit as NA increases. On the other hand, the length of the Gaussian beam ( $\mathrm{FWHM}_y$) drops even more rapidly as NA increases

(Fig. S10c). As the result of decreasing thickness and length as NA increases, the effective intensity shown in Fig. S10d exhibits tremendous enhancement as NA increases. The phototoxicity associated with such a high intensity may be problematic for living cell experiments.

### Optical scheme of the lattice lightsheet microscope

The schematic of the optical system is shown in Fig. S11. The beam from a laser combiner equipped with 488 nm (300mW, Coherent Sapphire 488 nm 300-CW), 561 nm (200mW, Oxxius LMX-561S-200-COL-PP) lasers is expanded to a diameter of 4 mm by two lenses (8 mm FL/ Ø1/2", Thorlabs C240TME-A, 20 mm FL/ Ø1/2" Edmund 47-661). The exposure time and the wavelength selection can be controlled by an acousto-optic tunable filter (AA Quanta Tech, Optoelectronic AOTF AOTFnC-400.650-TN) (**1**).

A pair of cylindrical lenses (Edmund NT68-160, 25 mm FL/12.5 mm dia (**2**); Thorlabs, ACY254-250-A (**3**)) is used to expand the beam in x axial direction. The expanded beam then passes through a polarizing beam splitter cube (PBS, Newport, 10FC16PB.3) (**4**) and a half-wave plate (Bolder Vision Optik, BVO AHWP3) (**5**), and uniformly illuminates on the central region of the spatial light modulator (SLM). The SLM itself consists of 2048 × 1536 ferroelectric liquid crystal pixels (Forth Dimension, QXGA-3DM) (**6**), which can change the polarity of the diffracted beam depending on the state of each pixel. The polarized beam can be imaged onto a custom quartz mask (**8**) by a polarizing beam splitter (**4**) cube and a lens (Edmund, 350mm FL / 50mm dia, VIS-NIR coating, achromatic lens (**7**)).

The lens pair (Thorlabs, AC254-100-A (**9**) and AC254-075-A (**10**) Ø1" Achromat, 400 - 750 nm) can reduce and image the beam from the mask to combine with Z axial galvanometer scanner (**11**). A relay lens (Thorlabs, AC254-85-A (**12** and **13**) Ø1" Achromat, 400 - 750 nm) combines two galvanometer scanners in the Z-axis (**11**) and the X-axis (**14**). After passing through two-dimensional scanning mirror sets, the beam is magnified through a relay lens (Thorlabs, AC254-254-A (**15**) and AC254-400-A (**16**) Ø1" Achromat, 400 - 750 nm) and conjugated to the back focal plane of the excitation objective (Special Optics, 0.66 NA, 3.74 mm WD) (**17**). The beam is projected onto the back focal plane of the excitation objective, and a self-reconstructed lattice beam is formed by optical interference at an incident angle of 32.8 degrees to the coverslip. Orthogonal to the illumination plane, water immersed objective lens (Nikon, CFI Apo LWD 25XW, 1.1 NA, 2 mm WD) (**18**) mounted on a piezo scanner (Physik Instrumente, P-726 PIFOC) (**19**) is used to collect the fluorescence signal, which is then imaged through an emission filter (Semrock Filter: FF01-523/610-25 and FF01-446/523/600/677) onto an sCMOS camera (Hamamatsu, Orca Flash 4.0 v2 sCOMS) (**21**) by a 500 mm tube lens (Edmund 49-290, 500 mm FL/50 mm dia; Tube Lens/TL) (**20**).

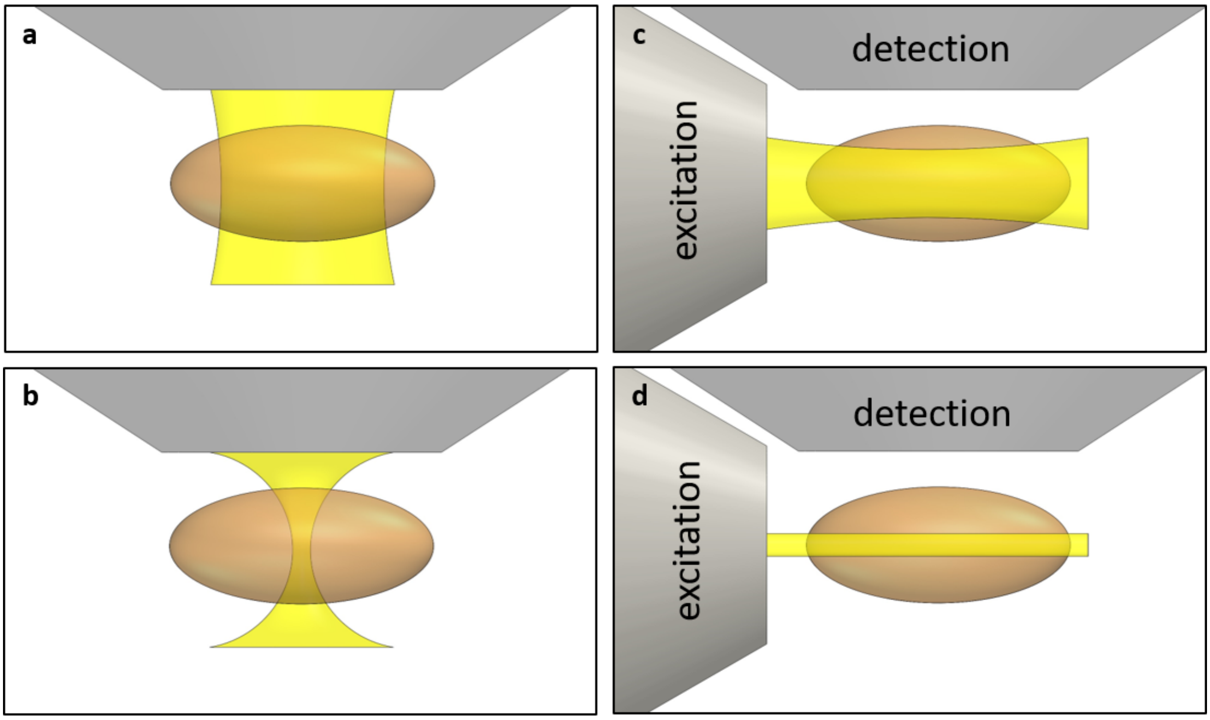


**Figure S1.** (a) Wide-field fluorescence microscopy employs a low NA objective to form a Gaussian beam excitation where the same objective is used for excitation and detection. In this case, there is no optical sectioning capability. (b) A high NA objective is used in confocal fluorescence microscopy with a Gaussian excitation beam where the same objective is for excitation and detection, and an additional pinhole is used to reject the out-of-focus background, providing optical sectioning capability. (c) A Gaussian beam is used in lightsheet microscopy, where a thick optical plane and a large field of view are used to confine the illumination at the part of the sample. A separate detection objective orthogonal to the excitation objective is used. (d) A lightsheet microscope employs a Bessel beam with a thin optical plane and a large field-of-view for confining the illumination at the specific part of the sample. A separate detection objective orthogonal to the excitation is used. A better optical sectioning capability can be achieved in Bessel beam lightsheet microscopy in comparison with Gaussian lightsheet microscopy.

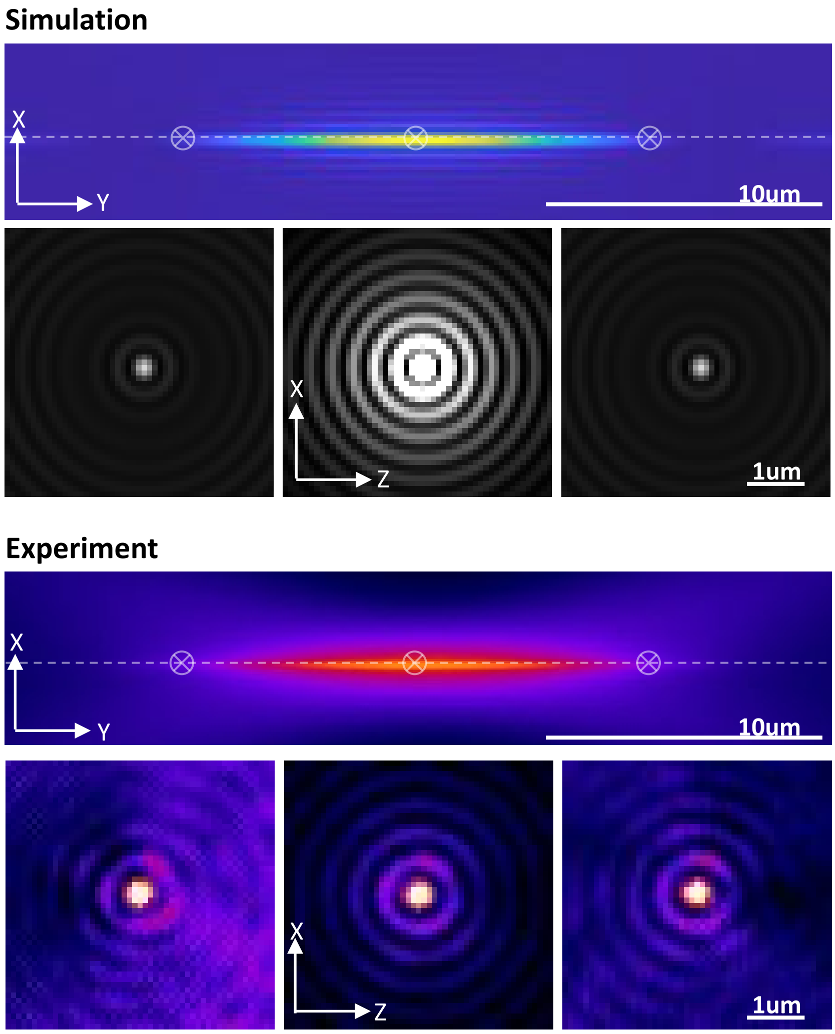


**Figure S2.** The calculated and observed energy distributions of a Bessel beam in XY and XZ planes, the energy distribution of the Bessel beam using the maximum and minimum numerical apertures (NA) of 0.64 and 0.56, respectively. The experimental Bessel beam profile was obtained by measuring the intensity profile of 100 nm fluorescent beads.

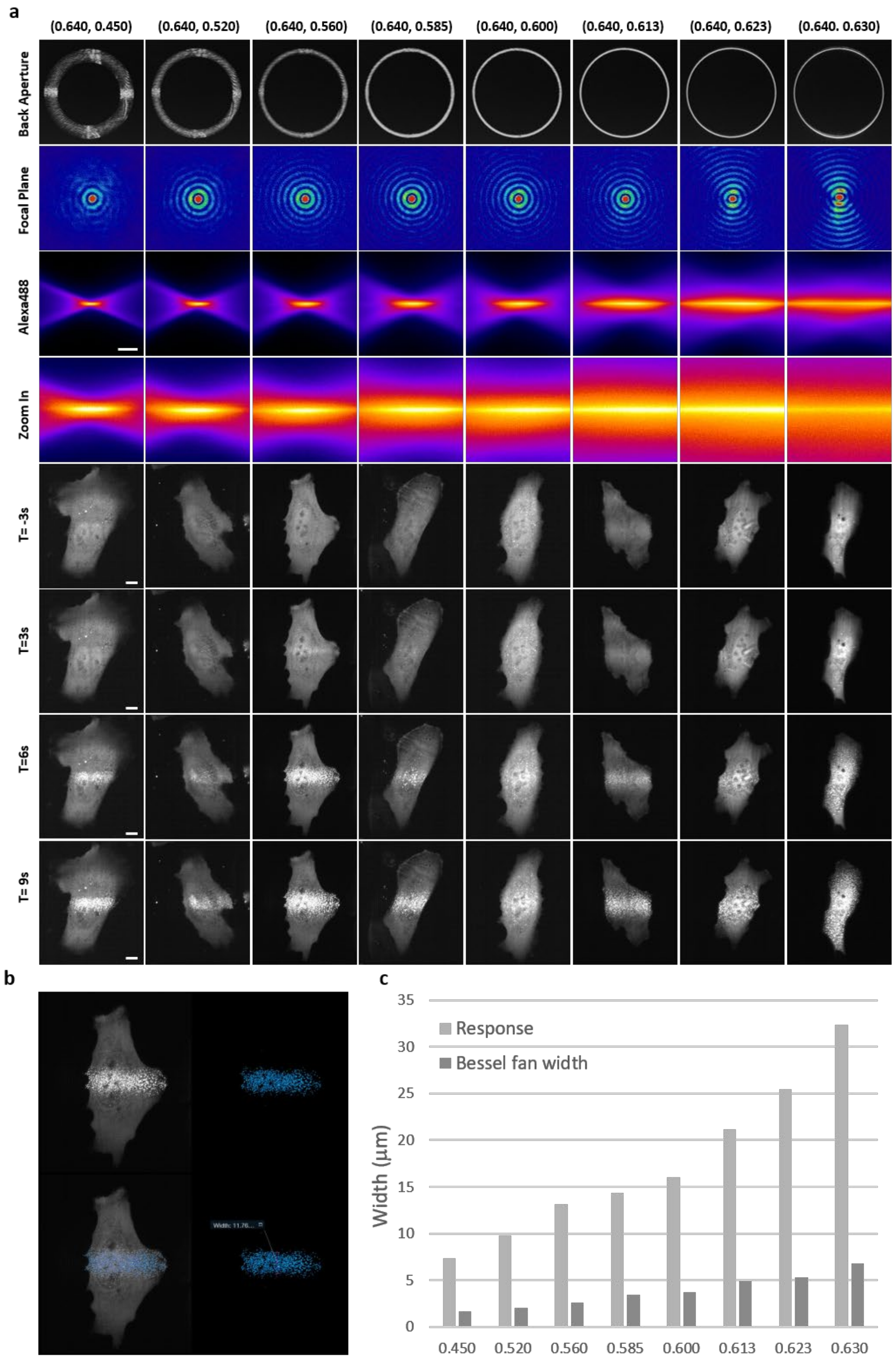


**Figure S3.** (a)The energy distribution in the XZ plane and the XY plane of the Bessel beam formed by a fixed maximum NA 0.64 and different minimum NAs (rows 1–4). Row 4 is a 4-fold enlarged view of row 3. Rows 5–8 are the corresponding time-lapse maximum intensity projection (MIP) images of a cell expressing CRY2olig-mRuby3 activated by various stimulation Bessel fans. (b) The processed images using the thresholding function in Amira. Top left: raw data, top right: processed result, bottom left: extracted subvolume in the central area, bottom right: view on X'Z plane. The width of optically induced clustering was calculated from the processed images. (c) The measured width of the induced clustering area (light gray) and calculated width (dark gray) of the stimulated beams. Scale bar 10 μm.

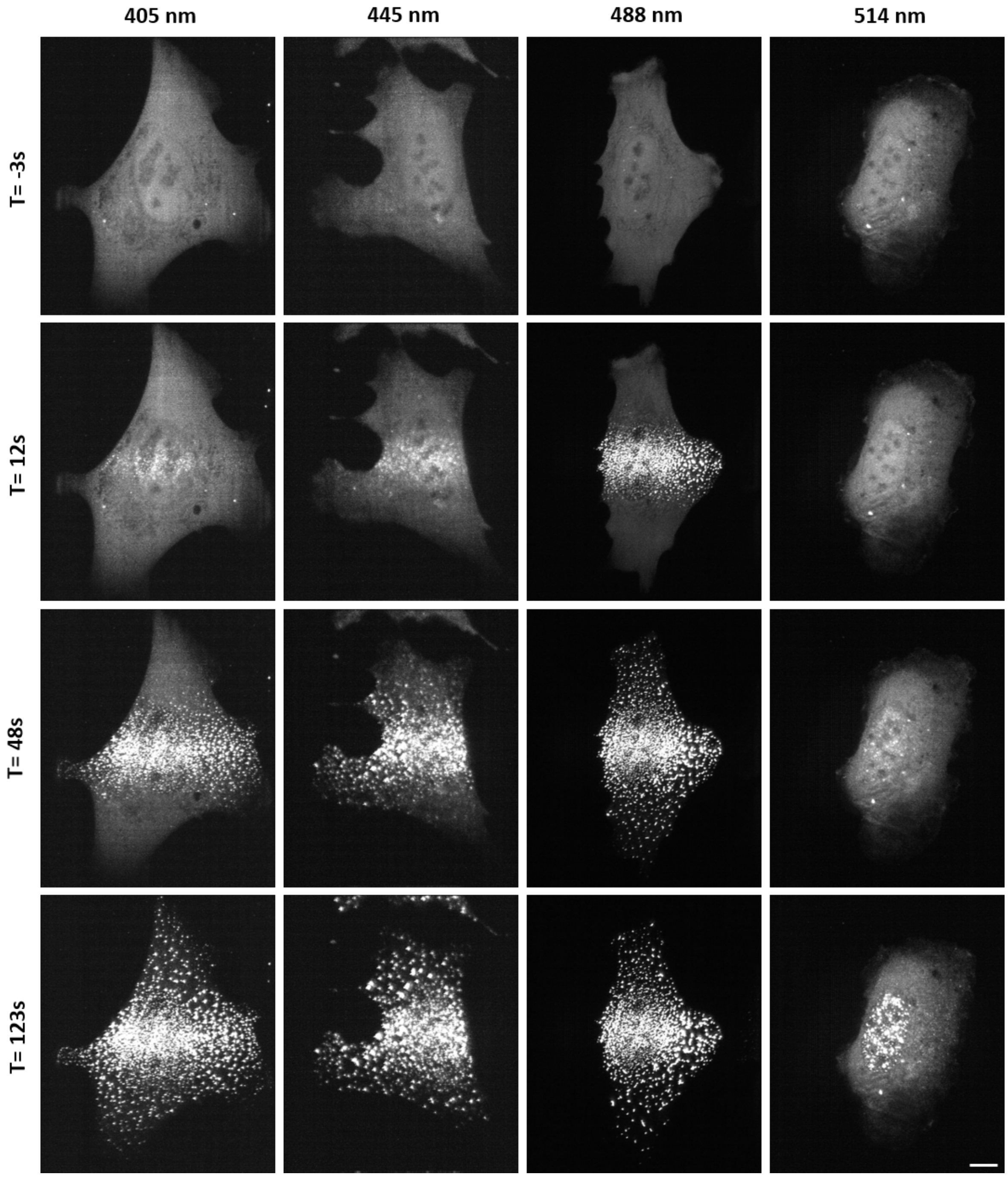


**Figure S4.** The time-lapse MIP images of cells expressing CRY2olig-mRuby3 stimulation by a different wavelength of Bessel fan at 1 nW. Scale bar 10 μm

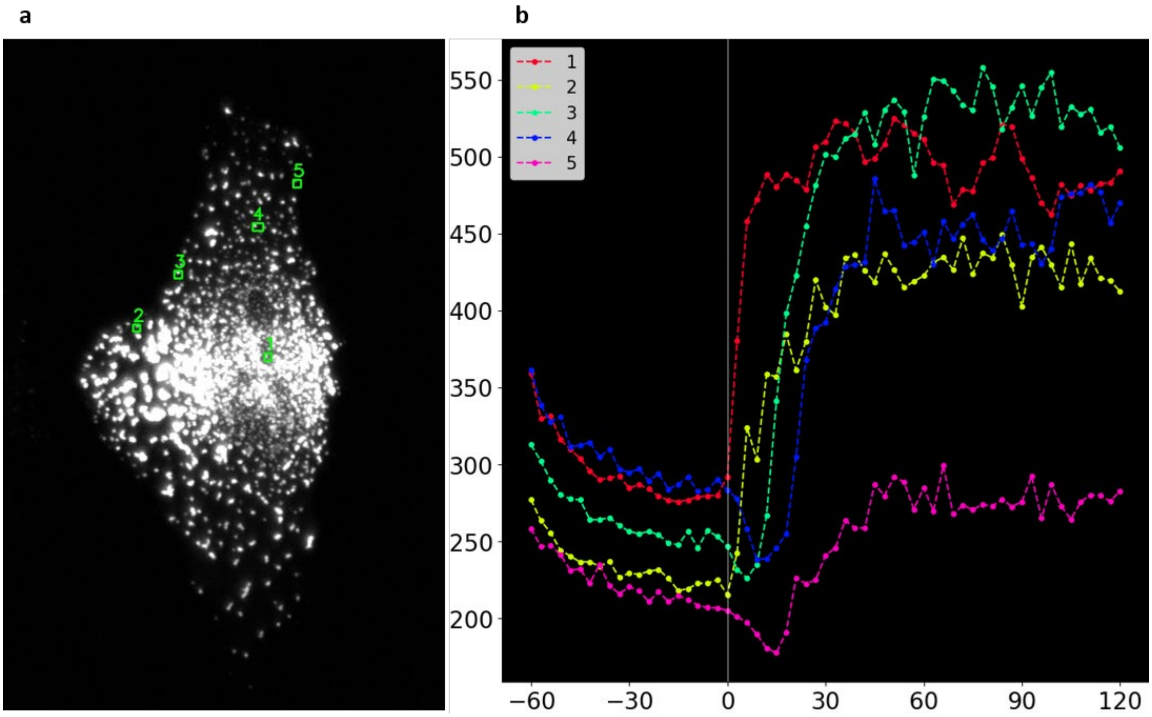


**Figure S5.** Characterization of the spatiotemporal behavior of the photoactivated clusters of CRY2oligo-mRuby3 expressed in the cell (a) illuminated by Bessel fan photoactivation schemes. (b) the time-dependent fluorescence intensities for the clusters marked in (a), the unit for the x-axis is second.

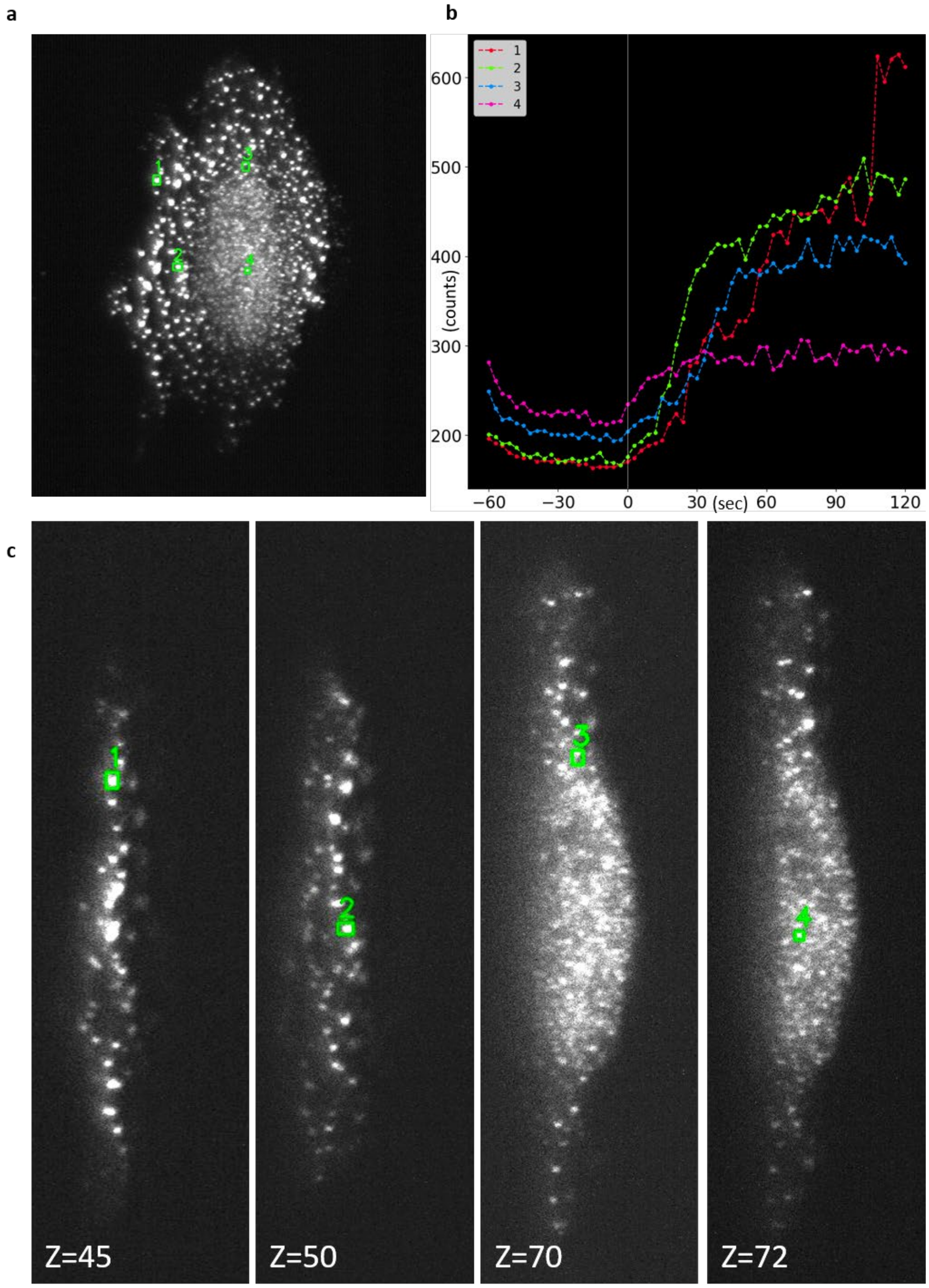


**Figure S6**. Characterization of the spatiotemporal behavior of the photoactivated clusters of CRY2oligo-mRuby3 expressed in the cell (a) illuminated by shifted Bessel fan photoactivation schemes. (b) the time-dependent fluorescence intensities for the clusters marked in (a). (c) the z slice images of the locations for the marked clusters among the 131 slices with z interval of 0.6 μm.

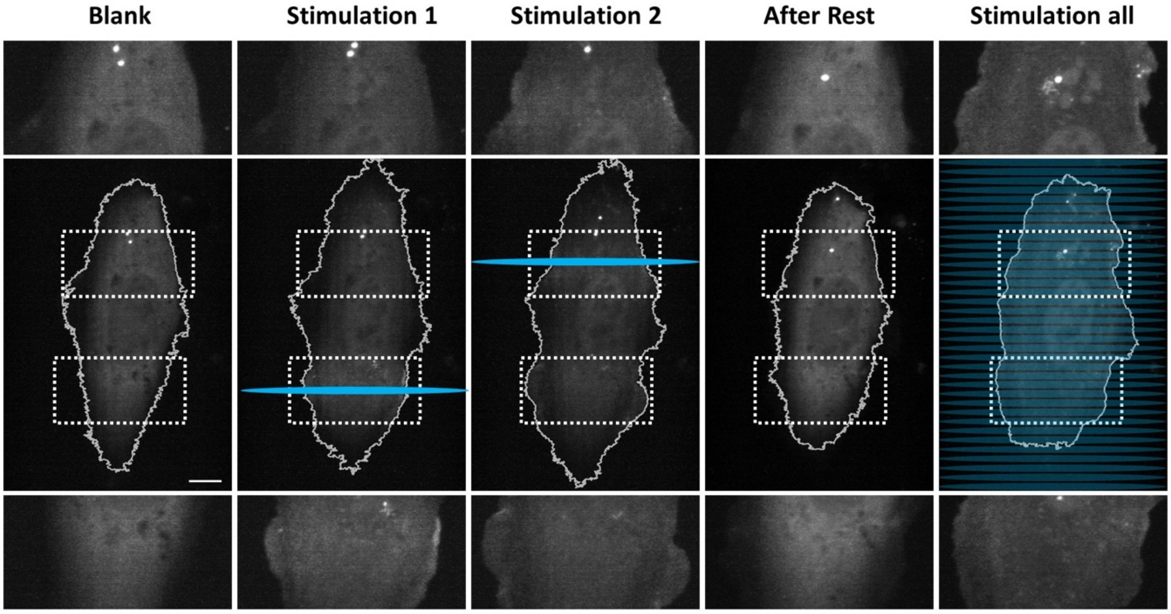
Blank
Stimulation 1
Stimulation 2
After Rest
Stimulation all

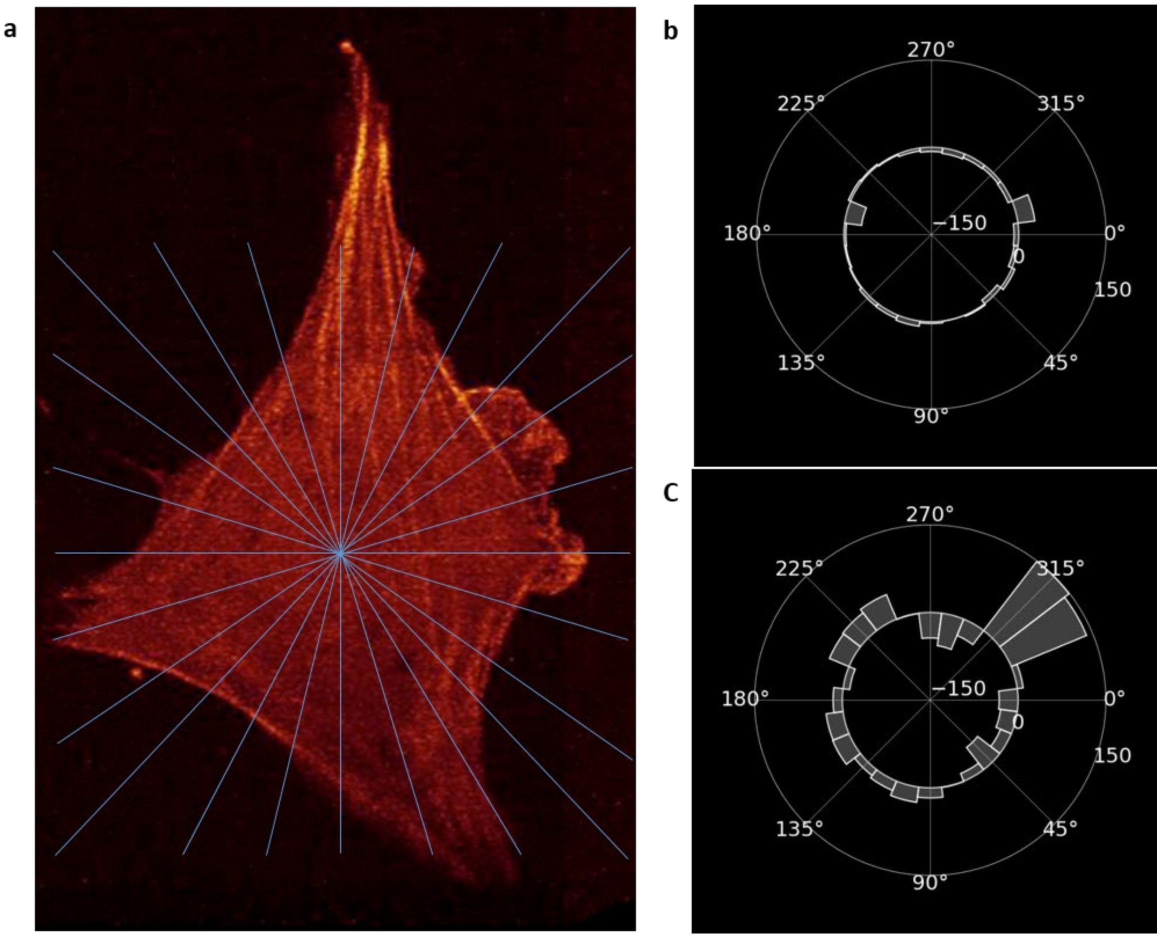


**Figure S8.** Quantification of the membrane protrusion and retraction during the guided cell migration. A polar coordinate is used to quantify the amount of protrusion and retraction at different parts of the cell with respect to the geometric center of the cell before and after photoactivation.

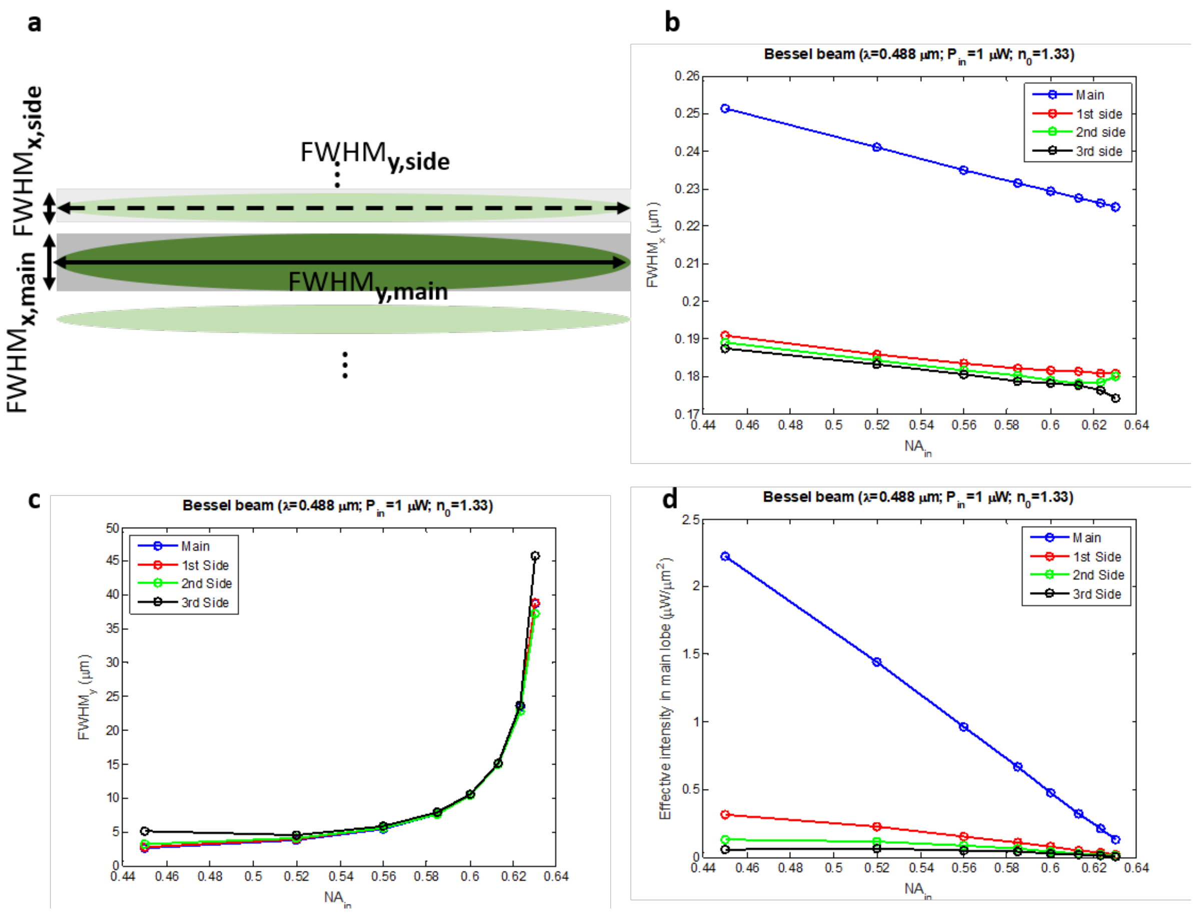


**Figure S9.** (a) Schematic of a Bessel beam profile at the excitation focus showing the beam length and beam thickness (b) The thickness of the main and side lobes of a Bessel beam with different inner NAs at a fixed outer NA= 0.64 at the wavelength of 488 nm and water environment (c) The length of the main and side lobes of a Bessel beam with different inner NAs at a fixed outer NA= 0.64 (d) The effective intensity of the main and side lobes of the calculated Bessel beams at an input power of 1μW.

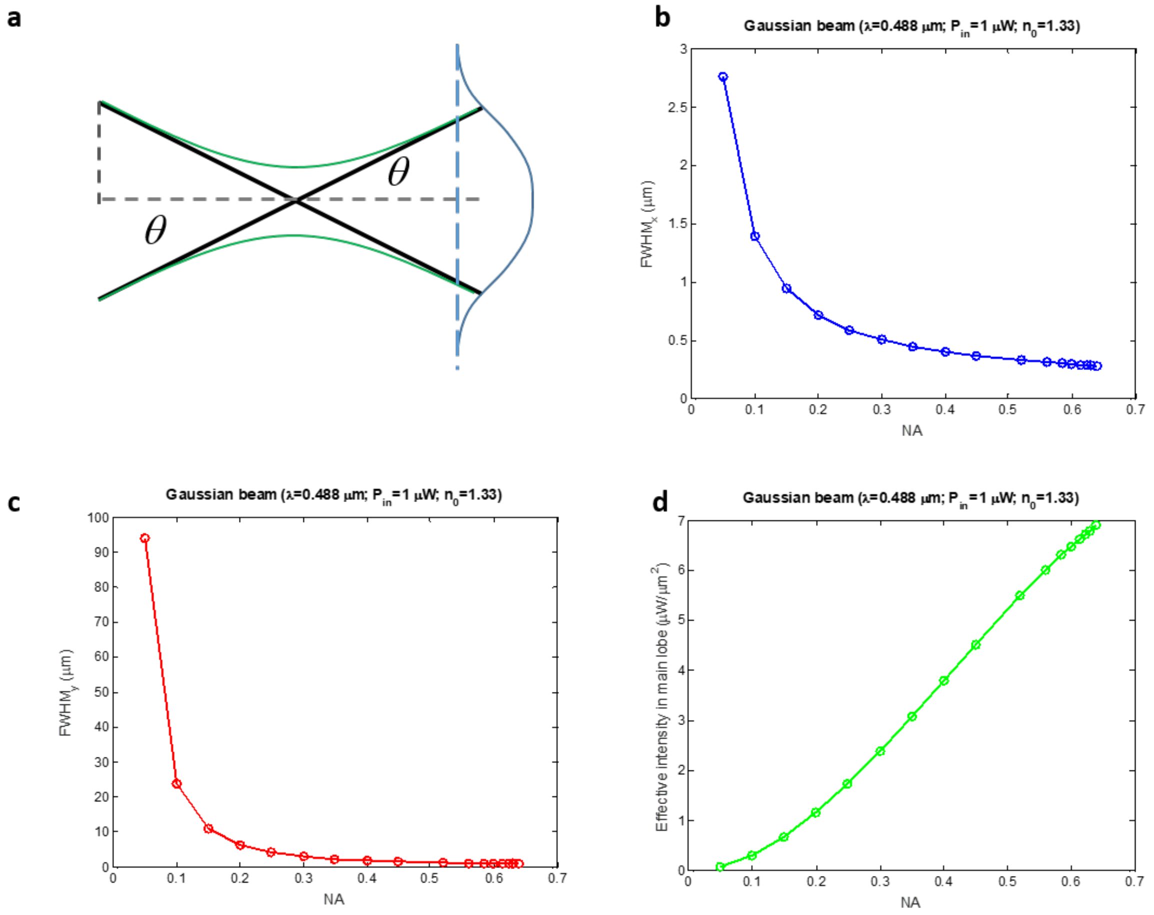


**Figure S10.** (a) Schematic of a Gaussian beam profile at the excitation focus showing the beam length and beam thickness (b) The thickness of a Gaussian beam with different excitation NAs at a wavelength of 488 nm and water environment (c)The length of a Gaussian beam with different excitation NAs (d) The effective intensity of the calculated Gaussian beams at an input power of 1μW.

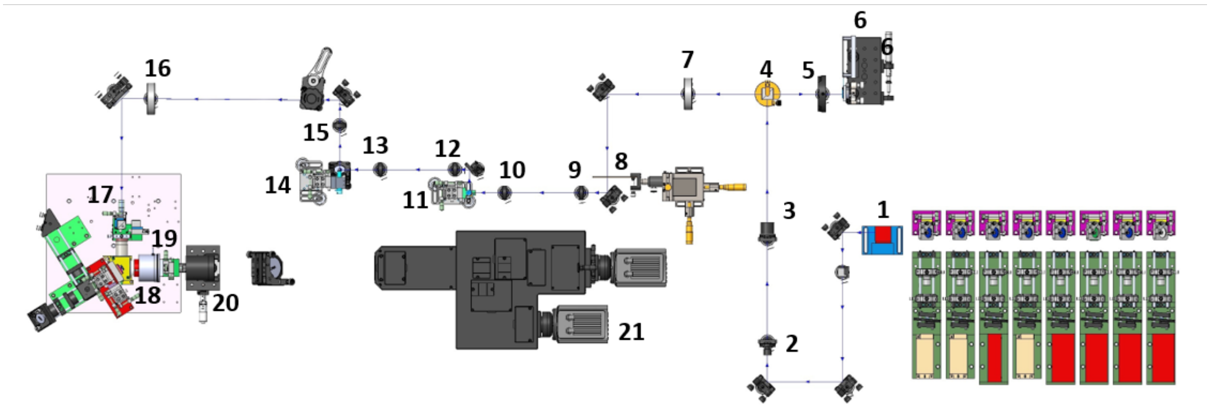


**Figure S11**. Optical scheme of the lattice lightsheet microscope. Each laser depends on its output beam profile to select the second lens of collimation for the equally expanded diameter of the beam (first lens: F= 8 mm; second lens: F= 20~30 mm). These collimated and expanded laser beams use dichroic mirrors (LM01-427, LM01-466, LM01-503, LM01-552, Do02-R561, and LM01-613) to combine and align to form a single beam passing through the acoustic optical tunable filter (AOTF). The laser beam selected by AOTF is extended in x-axial direction by a pair of cylindrical lenses (F= 25 mm and F= 250 mm) to fully cover the longer axis of the spatial light modulator (SLM). The x-axial direction extended laser beam is reflected by a polarization beam splitter (PBS) to project on SLM, where the turn-on area of SLM will transform the polarization of laser to form a patterned laser beam (lattice lightsheet). The patterned beam can directly pass throughout the PBS and a major focusing lens (F= 350 mm) to image on the Mask. For the spatial movement manipulation, the size of the lattice lightsheet is reduced to fit the size of galvo-mirror by a paired lens (F= 100 mm and F= 75 mm). At the same time, the z-axial and x-axial galvo-mirror's motion control need to balance by a paired lens (twin F= 85mm) connected to each other. After passing through SLM and two axial manipulations, the direction, height, and magnification of the lattice lightsheet beam are corrected by different optics (RS99 and F= 400 mm). The corrected lattice lightsheet will form on the front side of the excitation objective lens (special optic) and be imaged by the detect objective lens (Nikon 25X NA 1.1). Finally, the tube lens (F= 500 mm) is used to image the emission signals on the chip of sCOMs camera.

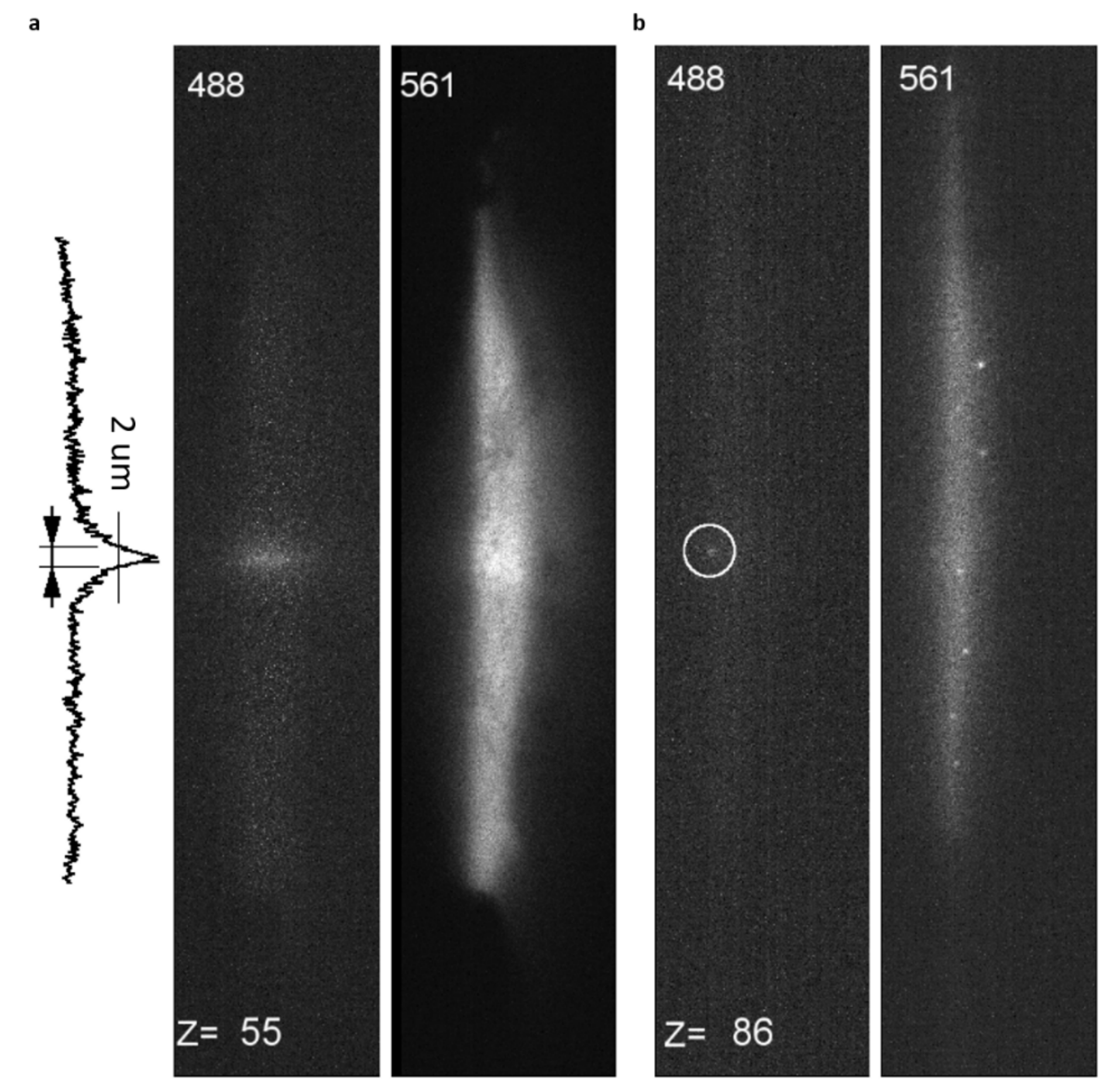


**Figure S12.** To characterize the Bessel fan and single Bessel beam activation, we measured the fluorescence signals of the photoactivated CRY2oligo-mRuby3 molecules by 488 nm stimulation. The weak fluoresce signals excited by 488 nm can be used to visualize the activated CRY2oligo-mRuby3 molecules within the path of the photoactivation beam, which could be used to indicate the beam profile of the photoactivation beam. In the Bessel fan activation (a), a strip with 2 μm width (FWHM) was observed from the contributions of main and side lobes of the illuminated Bessel beam at the z=55 out of 131 z-stacks. Note that all 131 layers are illuminated by the strip of the Bessel beam with a z interval of 0.6 μm. (b) A weak spot with a diameter of 2 μm was measured for single Bessel photo-activation at the z slice = 86, where only the selected plane was illuminated by the stimulation beam.

### Supplementary movie description

#### Movie 1

Time-lapse XY' MIP images of CRY2olig-mRuby3 expressed cells with various activation energies (0.5, 1, 2, and 4 nW). The interval between each stack is 3 seconds. The first stimulation starts at stack 20. The selected time point MIP images are shown in Fig. 1e. The video playback speed is 10 Hz.

#### Movie 2

Time-lapse XY' MIP images of CRY2olig-mRuby3 expressed cells stimulated with different wavelengths (405, 445, 488, and 514 nm) at 1 nW. The interval between each stack is 3 seconds. The first stimulation starts at stack 20. The selected time point MIP images are shown in Fig. S3. The video playback speed is 10 Hz.

#### Movie 3

Time-lapse images of optically induced clustering of CRY2oligo-mRuby3 in XY' Y'Z' XZ' planes by Bessel fan activation. The depth of the oligomerization is color-coded. The interval between each stack is 3 seconds. And the stimulation starts at stack 20. The selected images at different time points are displayed in Fig. 2b. The video playback speed is 10 Hz.

#### Movie 4

Time-lapse images of optically induced clustering of CRY2oligo-mRuby3 in XY' Y'Z' XZ' planes by shifted Bessel beam activation. The depth of the oligomerization is color-coded. The interval between each stack is 3 seconds. The first stimulation starts at stack 20. The selected images at different time points are displayed in Fig. 2b. The video playback speed is 10 Hz.

#### Movie 5

Time-lapse images of optically induced clustering of CRY2oligo-mRuby3 in XY' Y'Z' XZ' planes by single Bessel beam activation. And the depth of the oligomerization is color-coded. The interval between each stack is 3 seconds. The first stimulation starts at stack 20. The selected images at different time points are displayed in Fig. 2c. The video playback speed is 10 Hz.

#### Movie 6

Time-lapse XY MIP images of subcellular activation of the cell expressing CRY2mCherryiSH-p2a-CIBNcaax by Bessel fan activation, where the cyan color is the activation site. The interval between each stack is 3 seconds. The interval between each stack is 3 seconds. The first stimulation starts at stack 30 and restarts at stack 102 (rest time = 30min). The selected images at different time points are displayed in Fig. S4. The video playback speed is 10 Hz.

### Movie 7

Time-lapse XY' MIP images of optically induced membrane ruffling of cell expressing F-tractin-mCherry-p2a-CRY2iSH-p2a-CIBNcaax by single Bessel beam activation, where the cross is the activation site. The interval between each stack is 6 seconds. The stimulation starts at stack 30. The selected images at different time points are shown in Fig. 3a. The video playback speed is 10 Hz.

### Movie 8

Time-lapse XY' MIP images of optically induced cell migration of cell expressing EGFR-CRY2Olig-mApplex3 by Bessel fan activation, where the cyan line is the activation site. The interval between each stack is 6 seconds. And the stimulation starts at stack 20. The selected images at different time points are shown in Fig. 3b. The video playback speed is 10 Hz.

#### Movie 9

Raw 3D MIP image of optically induced cell migration of cell expressing EGFR-CRY2Olig-mApplex3 by Bessel fan activation. The cell was activated at T=0, Stack=60.The total stack number is 463 for about 6 hr observation. During the acquisition, the migrating cell was positioned at the imaging center of the field of view.

### Movie 10

The selected 18 XY' MIP images of cell migration for a cell expressing EGFR-CRY2OligI-mApplex3 activated by the Bessel fan. The cyan line represents the activation site. Different color codes are used to represent the contours of the cells at different time points. Selected time points are shown in Figure 4.

### Movie 11

The Raw 2D images used to reconstruct 3D volumetric image in the Bessel fan photoactivation experiment, where CRY2oligo-mRuby3 molecules are stimulated by 488 nm across the whole cell. 488 nm can excite the activated CRY2oligo-mRuby3 clusters with low efficiency, where the weak fluoresce signals reveal the contributions of main and side lobes of the illuminated Bessel beam forming a stipe of a width of 2 μm (left) as opposed to a lattice sheet imaging by 560 nm (right). Note that the residual electronic signal in the sCMOS leaking from the 560 nm channel was observed at the 488 nm channel.

### Movie 12

The Raw 2D images used to reconstruct 3D volumetric image of a single Bessel beam photoactivation experiment. Only a weak fluorescent signal was observed at the selected plane (z=55), where the cell was illuminated by 488 nm (left). In comparison, a lattice light-sheet imaging at 560 nm was used to record the fluorescence signals of the cell (right).
